# Modeling mucociliary mixing and transport at tissue scale

**DOI:** 10.1101/2025.01.06.631467

**Authors:** Ling Xu, Pejman Senaei, Yi Jiang

## Abstract

Mucociliary clearance is the primary defense mechanism in our respiratory system against aerosol pathogens and allergens. The rhythmic movement of cilia on airway-lining cells propels mucus flow, driving the movement of trapped particles. We want harmful particles, e.g., pathogens and allergens to be transported rapidly out of the airway and beneficial particles, e.g. vaccines and medicine, to stay and disperse. However, the impact of cilia density and distribution on mucociliary mixing and transport at the tissue scale remains poorly understood. We present three-dimensional (3D) simulations of mucociliary mixing and transport at the tissue scale. We investigate the influence of ciliary cluster spacing, metachronal wave, and ciliary density on mucus mixing and transport. Our findings reveal that: (i) cilia clusters produce swirls of mucus flow, their size scales with ciliary density; (ii) a single cilia cluster generates horizontal and upward transport with horizontal mixing; (iii) for three aligned cilia clusters, an optimal cluster spacing exists for achieving maximum horizontal transport; (iv) asynchronous beating among clusters enhances mixing but hinders transport; and (v) the diffusion of particles exhibits spatial inhomogeneity, leading to particle aggregation into discrete groups over time. These discoveries have implications for understanding particle distributions in the airway and contribute to drug delivery design considerations.

**Author summary:** Our airways are exposed to various foreign particles, including allergens, pathogens, and aerosolized vaccines or medications. The mucus coating the airway surface traps these particles, and the beating cilia extending from the ciliated epithelial cells play a crucial role in propelling the flow of mucus to transport the particles out of the airway. This mucociliary clearance process is closely associated with the initiation and progression of airway tissue response, such as allergic responses, infections, and treatment effects. Changes in ciliated cell density or mucus production can significantly impact this process. Conversely, achieving a homogeneous distribution of aerosol drug particles within the airway is essential for optimizing treatment outcomes. We simulate cilia-mucus coupled flow to examine how the motion of cilia influences the transport and mixing of particles in the mucus. Our key findings suggest that an optimal spacing for ciliated cell clusters that enhances directional transport. Intriguingly, while the metachronal wave promotes mixing, it also tends to hinder the overall transport of particles.

## Introduction

Mucociliary clearance serves as the primary defense mechanism of the airways, playing a crucial role in expelling foreign particles. The airways are lined with a mucus layer that effectively traps inhaled particulates. Beneath the mucus, the cilia, tiny hair-like structures protruding from the airway epithelium, beat rhythmically. This coordinated movement propels mucus, along with trapped particles, out of the airway, contributing to the respiratory defense system [1]. Conversely, when administering nasal sprays for vaccines [2] or treating allergic rhinitis [3], the goal shifts towards achieving maximal mixing with minimal mucus transport.

The airway epithelium primarily consists of the ciliated cells where cilia reside, the goblet cells responsible for mucus secretion, and the basal cells, which can proliferate and differentiate to maintain epithelial integrity. Different segments of the airway exhibit distinct cellular compositions [4]. A microscopy image illustrating the distribution of cilia in the rabbit trachea (Fig. 1a) provides insight into their arrangement. A typical cilia cluster has a dimension of 10*µ*m and comprises 100-200 cilia on its apical surface [5] (Fig. 1b). Each cilium is anchored at its base and undergoes periodic beating in a three-dimensional (3D) circular motion, including an effective stroke and a recovery stroke. During the effective stroke, the cilium remains rigid and upright, while it becomes pliable and bends during the recovery stroke [6]. It is well-established that the coordinated beating of cilia generates a net unidirectional flow of mucus, effectively transporting foreign particles or pathogens out of the airway.

**Fig 1.**
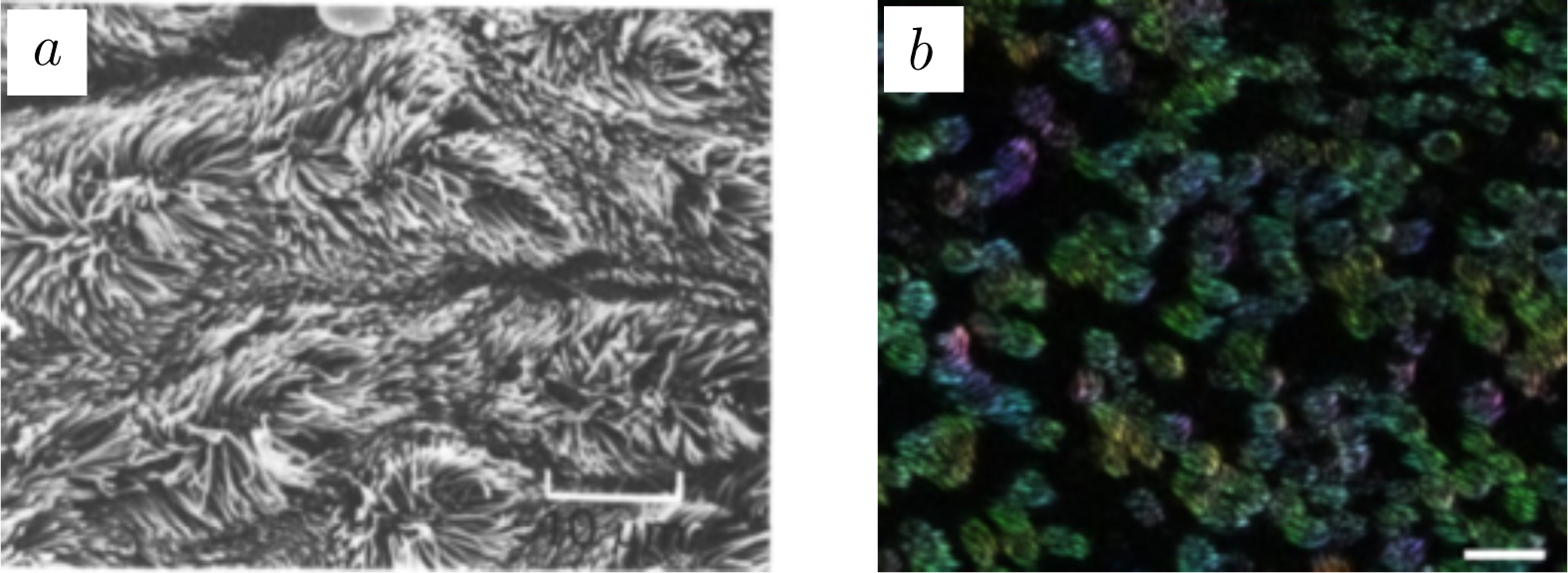
Tissue-scale cilia images.(a) Cilia in the rabbit tracheal from Sanderson and Sleight [6]. Scale bar represents 10*µ*m. (b) Cilia orientation map in human bronchial cultures obtained using over 500 images, from Khelloufi et al. [5]. Scale bar represents 20*µ*m.

Several airway diseases, such as primary dyskinesia, chronic obstructive pulmonary disease (COPD), and asthma, can impair the ciliary function [7]. Using human bronchial cultures ranging in size from micrometers to centimeters, Khelloufi et al. found, underneath the mucus, a circular order of the ciliary beating directions (swirl) drives the mucociliary transport and the swirl size was shown to scale with ciliary density [5]. Gsell et al. further demonstrated, using human bronchial epithelium reconstituted in-vitro and a 2D hydrodynamic model, that increasing ciliary density increases the swirl size, leading to longer-range unidirectional mucus flow [8].

Many respiratory viruses, including influenza virus and SARS-CoV-2, target the ciliated cells [9–13]. While the lung epithelium can sustain normal function under mild injuries, permanent damage may occur if large areas of ciliated cells die due to infection [4]. SARS-CoV-2 infection, in particular, can inflict severe damage on the lung epithelium, resulting in serious pneumonia, acute respiratory distress, and even death [13, 14]. A recent model for within-host SARS-CoV-2 infection incorporates mucociliary clearance as a crucial mechanism in viral transmission [15]. However, how ciliary density change as infection progresses and how that impacts mucociliary transport has not been studied.

The mathematical exploration of the mucociliary system has spanned over six decades since Taylor’s work in 1951 [16]. Early investigations focused on the wavy shape of cilium-like organelles, modeling them as sinusoidal functions [17]. Brokaw proposed the shear mechanism involving internal doublet structures to elucidate the undulating motion [18]. Lighthill introduced the slender body theory [19] that represents the beating cilium body as a line of singular forces resulting from the self-activation and self-regulation of the motile body, and the fluid flow velocity near the motile body is proportional to the force exerted. Barton and Raynor made the assumption that feedback from the fluid flow is negligible [20]. They modeled the cilium as a rigid rod, straightening during the effective stroke and automatically shortening during the recovery stroke. Building upon the slender body theory, Cortez introduced the method of regularized stokeslets, designed to solve the Stokes equations in free space for zero Reynolds number flow [21]. This method has been widely adopted for simulating swimming micro-organelles and other biological fluid flow problems, where inertial forces are negligible [22]. For instance, the Reynolds number within the mammalian respiratory tract is approximately 0.01, where viscous forces play a dominant role in the mucociliary system [23]. Guo et al. explored the in-phase and anti-phase synchronization modes of two adjacent cilia [24]. They modeled cilia as chains of beads, each bead is driven by an external periodic force as a rower [25]. Furthermore, Charkrabarti and colleagues modeled a lattce of cilia as rowers to investigate the formation and maintenance of metachronal waves [26]. They findings suggest that spatial inhomogeneities contribute to the robustness of the metachronal waves, enhancing long-range transport.

Other mathematical models have delved into various facets of the mucociliary system. Osterman and Vilfan determined optimal beating patterns of cilia based on considerations of energetic efficiency [27]. Studies by Eloy and Lauga have examined the kinematics of the most efficient cilium [28]. Yang, Dillon, and Fauci explored the emergence of cilia beating synchrony through hydrodynamic effects [29], highlighting the complex interactions between fluid dynamics and ciliary activity. We sought to identify key factors influencing cilia motion and particle transport in mucociliary clearance [23]. The advent of supercomputing has enabled large-scale computational studies in this domain. Mitran’s work featured a full 3D simulation involving 256 beating cilia [30]. Elegeti and Gompper simulated 400 2D cilia arrays within a 3D fluid flow [31]. Additionally, Ma and Lutchen simulated aerosol deposition using a turbulence model [32]. Their sensitivity analysis highlighted the crucial dependence of simulation outcomes on the characteristics of the flow field [32]. These studies collectively underscore the importance of computational models in unraveling the nuances of mucociliary clearance and its implications for respiratory health.

The objective of this study is to address how ciliary density and distribution impact the mucociliary transport and mixing at the tissue level. We adopt the regularized stokeslet method, assuming the mucus flow to be a Stokes flow with infinite viscosity [33]. Extending from our previous work [23], we model the 3D cilium beating as a rigid rod with a prescribed motion. We vary the cilia cluster spacing and density to study their impact on the trajectories of massless particles placed in the flow. These particles not only help to illustrate material transport and mixing, but also mimic the motions of tiny particles such as viruses whose sizes are negligible.

We adopt the regularized stokeslet method, assuming the mucus flow to be a Stokes flow with infinite viscosity [33]. Instead of modeling cilium motion as rowers [20], we prescribe the 3D motion of all beating cilia. This approach allows us to investigate the effects of other parameters, including cilia density, cilia cluster spacing, and metachronality. The primary focus of our work is on the transport and mixing dynamics involving multiple cilia clusters at tissue scale. We vary the cluster spacing and tissue cell density to explore their impact. To illustrate material transport, we introduce massless particles into the fluid flow, mimicking the motions of bacteria or viruses. Our simulations unveil intriguing global transport phenomena and diverse local mixing patterns. Additionally, we examine the phase shifts of metachronal waves among cilia clusters, discovering that such shifts can hinder overall transport. Further details are provided in the numerical results section.

### A Tissue-scale 3D Mucociliary Flow Model

To simulate mucociliary flow in 3D at the tissue scale, we integrate a cilium beating model and a mucus flow model. We set up the computation model of cilia clusters, and vary the parameters corresponding to the ciliary density and distribution.

### Cilium beating model

We model the cilium as a rigid rod with a length *L* that changes during the forward and backward strokes in cilium beating motion [23]. Figure 2(a) illustrates the motion of the rod in one revolution. The position of rod ***x*** = (*x, y, z*) is

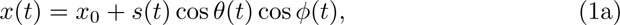

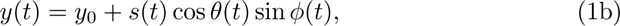

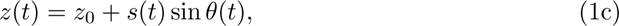

**Fig 2.**
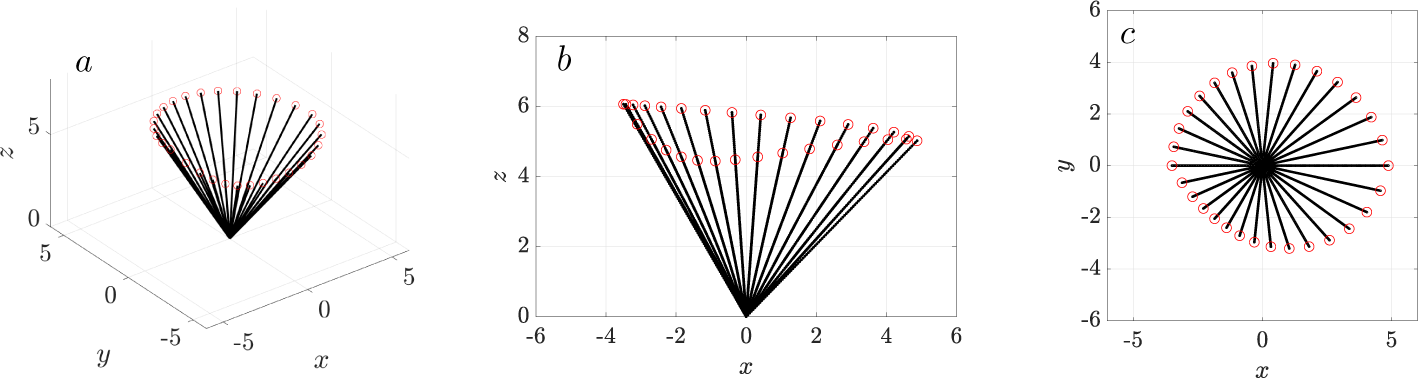
Illustration of a 3D beating cilium simplified as a rotating rod with varying length. (a) The rod’s motion over one full revolution, with red circles marking the rod tips. (b)Projection of the rod’s profile onto the (*x, z*)-plane, (c) Projection of the rod’s profile onto the (*x, y*)-plane.

***x***_0_ = (*x*_0_*, y*_0_*, z*_0_) is the base of the rod on the (*x, y*)-plane (*z*_0_ = 0), *s* ∈ [0*, L*] is the Lagrangian parameter. The instantaneous velocity of the rotating rod is computed analytically using ***u*** = d***x****/*d*t*.

Figures 2(b,c) show the rod’s profile projected onto the (*x, z*)- and (*x, y*)- planes. The rod rotates at a given angular velocity *ω* in the counter clockwise direction about *z*-axis. The azimuthal angle *ϕ* and the polar angle *θ* both time-dependent. The azimuthal angle is *ϕ*(*t*) = *ϕ*_0_ + *ωt*, *ϕ*_0_ is the initial phage lag of the rod at time *t* = 0. The polar angle is *θ*(*t*) = *θ*(*ϕ*(*t*)), and it describes the angle from the (*x, y*)-plane. The range of the polar angle *θ* is limited to [0.8, 2.4] radians based on the experimental depiction of a typical beating cycle of a cilium [6],.

We previously simplified the cilium as a rigid rod with a varying length *L*(*ϕ*) to mimic the forward (power) stroke and the backward (recovery) stroke [23]. We adopt the same description here:

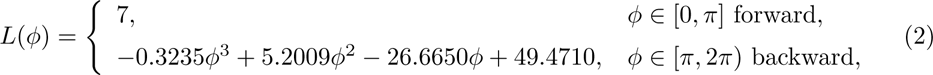

As shown in Fig. 2(c), the trajectory of the rod tip is not circular on the (*x, y*)-plane due to the variation of both *θ* and the rod length.

### Cilia-driven mucus flow

The mucus is modeled as an incompressible Newtonian fluid, where the inertial forces are negligible [34]. The governing equation is:

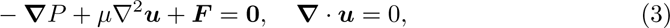

where *P* is the fluid pressure, ***u*** is the fluid velocity, *µ* is the fluid viscosity, and ***F*** (***x***) represents the force density exerted on the fluid by the cilia. Following the work of Guo et al. [24], we choose a regularized Stokeslet formulation [33] where forces are distributed along the rod. The force density at a point ***x****_c_* follows a distribution

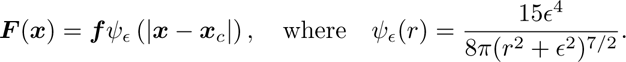

Here ***f*** is the force coefficient, *ψ_∈_*(*r*) is the regularized kernel, and *r* = |***x*** − ***x***_0_|. We choose *c* = 0.1, which corresponds to the cilium thickness. The fluid flow is subject to no-slip condition at the bottom wall or epithelial cell surface, ***u*** = **0** at *z* = 0. As the stiff rod beats, the force coefficients ***f*** of all stokeslets are updated according to the instantaneous rod velocity.

### Tissue-scale cilia clusters

Figure 3(a) illustrates an array of three cilia clusters, each measuring 8*µ*m × 8*µ*m in size. Within each cluster, 3 × 3 are arranged in a grid with equal spacing of *D* = 4*µ*m. The spacing between cilia clusters, denoted as *D_c_*, is variable in this study. The array is aligned along the *x*-direction, corresponding to the length of the airway, and transport dynamics along this direction is of primary interest. Because a single cliated cell in the airway epithelium contains approximately 100 to 200 cilia [35], each model cilium in our study can be considered as representing the synchronized motion of hundreds of cilia on a single ciliated cell.

**Fig 3.**
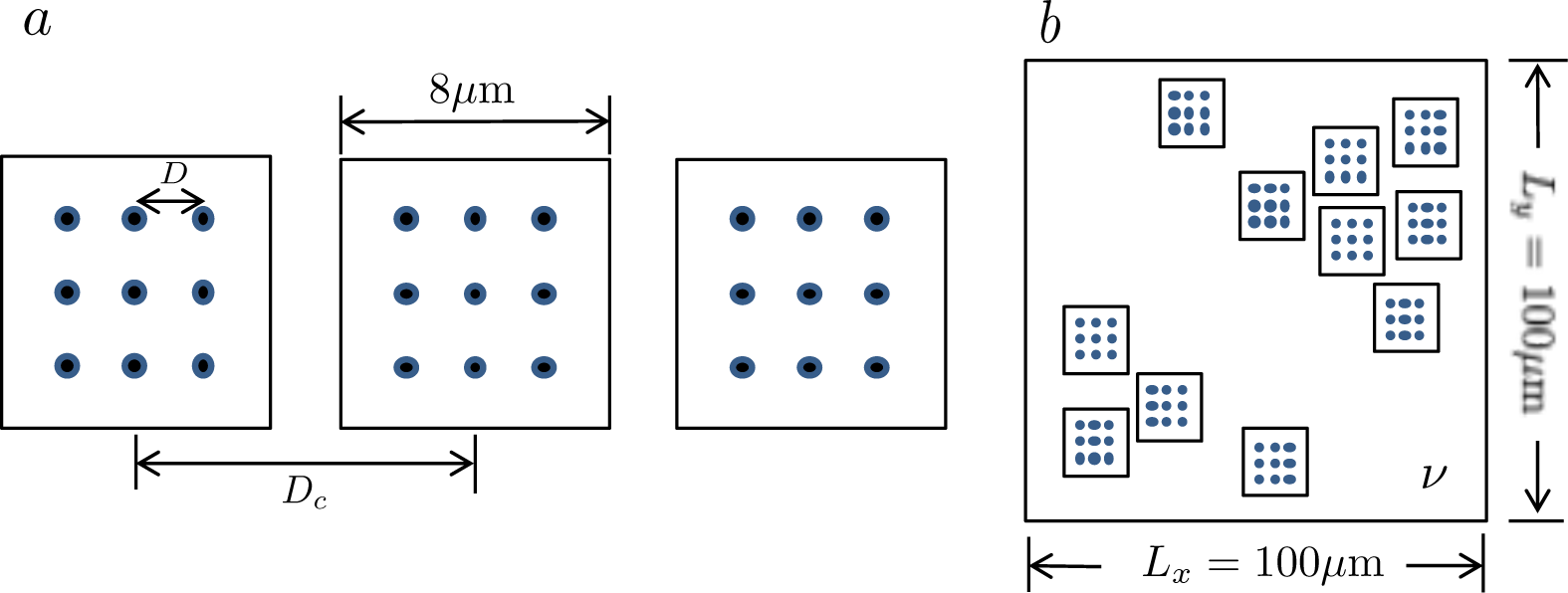
Illustration of the tissue-scale cilia cluster model. (a) The top view of an array of three cilia clusters, each containing nine equally spaced cilia. The spacing between individual cilia within a cluster is *D* = 4*µ*m, while the inter-cluster spacing is *D_c_* (variable). (b) A large patch containing multiple clusters, covering an area of 100*µm* × 100*µm*. *ν* is the variable ciliary density. Blue dots indicate the position of individual cilia.

Figure 3(b) illustrates a larger area of the epithelium containing multiple ciliated clusters. The area is represented as a square with side lengths of *L_x_*= *L_y_*= 100*µ*m. The ciliary density, *ν*, is the ratio of the total area occupied by the cilia clusters (small squares) to area of the larger square region. These cilia clusters are distributed randomly within this region. Table 1 lists all computational parameters used in this study.

**Table 1.** Parameters used in this study are based on experimental measurements.

| Parameter | Symbol | Dimensional value | Reference |
| --- | --- | --- | --- |
| Maximum rod length | $\max(L)$ | $7\mu\text{m}$ | [35] |
| Rod/Cilium beating frequency |  | 18 Hz | [35] |
| Rod/Cilium angular velocity | $\omega$ | $36\pi\text{s}^{-1}$ | |
| Rod spacing in the cilia cluster | $D$ | $4\mu\text{m}$ | |
| cilia cluster spacing | $D_c$ | $3D-12D$ | |
| Ciliary density | $\nu$ | 0.1, 0.2, 0.4 | [5] |
| Tissue patch | $L_x, L_y$ | $100\mu\text{m}$ | |

## Results

### Higher ciliary density leads to larger swirls and more directed transport

To investigate particle motion driven by mucociliary flow at the tissue scale, we simulate a patch of ciliated cells. The patch size is represented by a square region with dimensions spanning [−50, 50] × [−50, 50]*µ*m. Withi this patch, cilia clusters are randomly distributed with a density *ν*. The coordinated beating of these cilia drives mucus flow. At a low cilia density (*ν* = 0.1), the 3D velocity field of the cilia patch shows rotational order (Fig. 4), resulting in less predictable transport directions.

**Fig 4.**
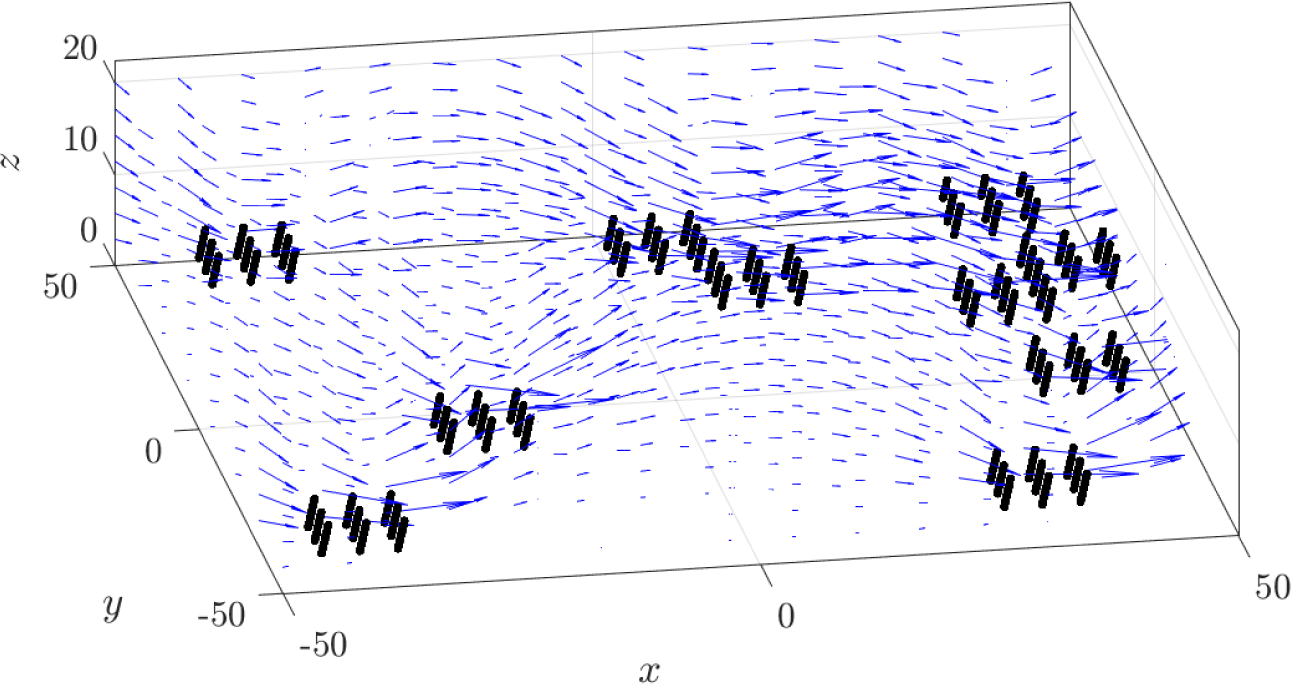
Typical snapshot of the swirl pattern in mucus flow driven by cilia at a density of *ν* = 0.1. Blue arrows indicate the velocity field. Black lines represent cilia, beating synchronously with an initial phage *ϕ*_0_ = 0.

To examine how the swirl motion drives particle distribution and its dependence on ciliary density, we compare mucociliary flow patterns at three cilary densities: *ν* = 0.1, 0.2, and 0.4. The ciliated cell density in human bronchial culture ranges from 0 to 0.70, depending on the location in the airway and pathological conditions, with an overall average of 0.15 [5]. Our density choices reflect this physiological range.

Fig. 5 shows the top view of particle distributions in a ciliated tissue patch, with time and time ciliary density. The massless particles display swirls as those in the human bronchial cultures (Ref. [5], figure 3B). With time, from top to bottom (Fig. 5), the initial well-segregated particles of different colors show the emergence of multiple counter-clock-wise swirls. The number of swirls is far fewer than the number of cilia clusters, suggesting that the swirls are the consequence of hydrodynamic effects of between the cilia clusters. The swirl size increases in time, with the bulk of the particles tending to the lower left of the simulation domain. At *t* = 11.11*s*, after 400 revolutions, the majority of the particles remain segregated. From left to right, the swirl size increases with ciliary density, consistent with the in vitro observations using human bronchial epithelium [8]. Higher ciliary density shows more pronounced mixing.

**Fig 5.**
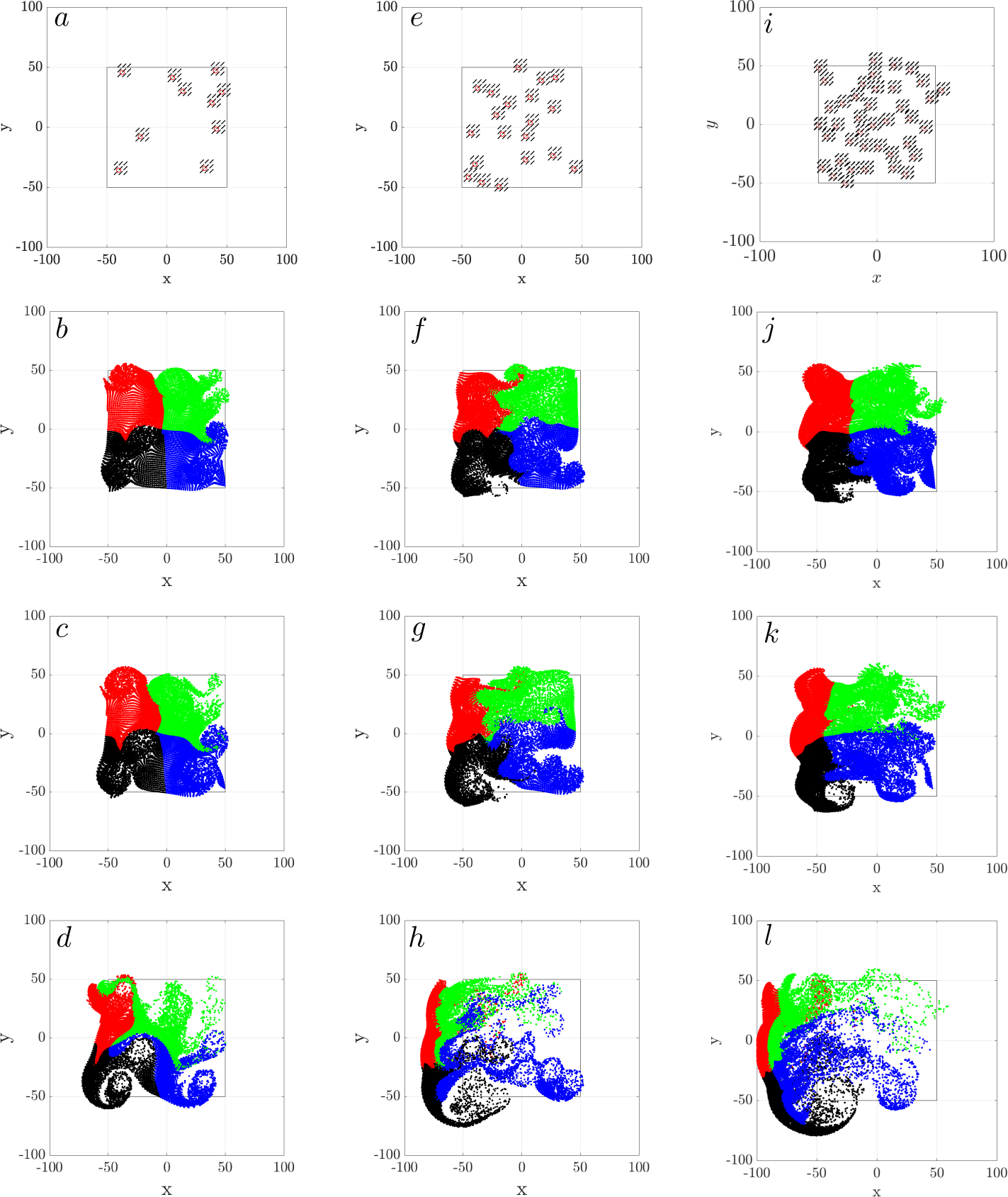
Swirl motion of passive particles as a function of ciliary density: (a-d) *ν* = 0.1 (e-h) *ν* = 0.2, (i-l) *ν* = 0.4. The top row (a, e, i) shows the locations of cilia clusters. The remaining rows show the top view of the distribution of passive particles at *t* = 1.38*s* (b, f, j), *t* = 2.78*s* (c,g,k), *t* = 11.11*s* (d, h, l). These time points correspond to 50, 100, and 400 cilia beating revolutions, respectively, with all cilia beating synchronously. Particles are initially distributed in four color-coded groups (black, blue, green, and red) within the ciliated region [−50, 50] × [−50, 50]*µ*m and *z* ∈ [5, 9]*µ*m. The simulation domain is [−100, 100] × [−100, 100]*µ*m.

We calculate the center of mass (CoM) of all particles to quantitatively compare transport efficiency as a function of cilary density (Figure 6). The results for each density value (*ν*) represent the average of three independent simulations. For all three densities, the net particle transport occurs in the negative *x*- and *y*-directions and the positive *z*-direction, indicating particles movement toward the lower-left corner of the simulation domain and away from the cell surface. Higher ciliary density results in faster and more extensive directed transport, consistent with in vitro observation [8].

**Fig 6.**
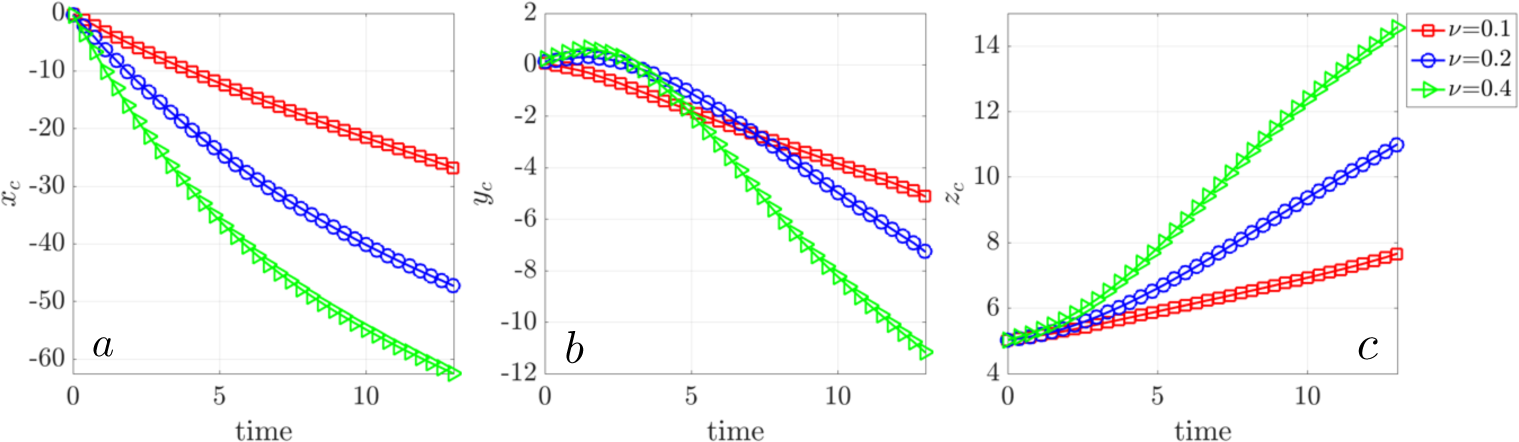
Net particle transport driven by mucociliary flow at ciliary densities *ν* = 0.1 (red), *ν* = 0.2 (blue), and *ν* = 0.4 (green). (a) Center of mass (CoM) displacement in the *x*-direction, (b) CoM displacement in the *y*-direction, (c) CoM displacement in the *z*-direction.

To better quantify the movement of the particles in mucociliary flow and analyze vertical mixing, we calculate the mean square displacement (MSD), defined as *MSD*(*τ*) = ((***z***(*t* + *τ*) − ***z***(*t*))^2^), for particles at various initial heights (Figure 7). A reference line of slope 1 is included to indicate the behavior of simple Brownian motion. Massless particles near the base of the cilia (*z* = 1) do not move much, hence their MSD is low across all ciliary densities. Particles away from the bottom of the mucus travel more as driven by the cilia. At low ciliary density (*ν* = 0.1), the MSD curves at all heights have slopes less that 1, indicating that particles move less than Brownian motion (subdiffusion). Increasing ciliary density leads to increased motion and larger MSD slope.

**Fig 7.**
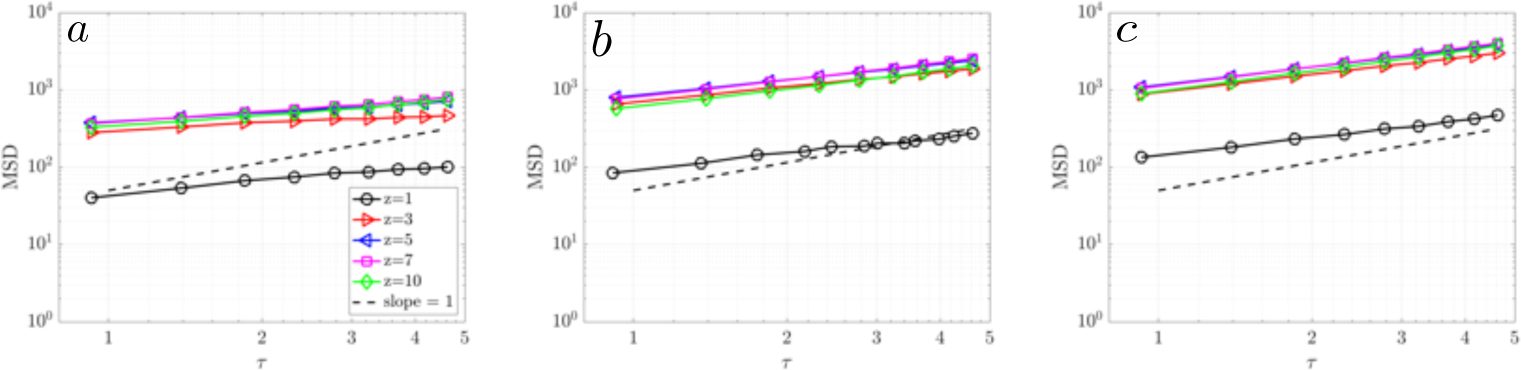
Mean square displacement (MSD) of massless particles at various initial vertical heights (*z* = 1, 3, 5, 7, 10*µ*m) for ciliary densities: (a) *ν* = 0.1, (b) *ν* = 0.2, (c) *ν* = 0.4. The gray dashed reference line indicates a slope of 1.

To quantify particle aggregation and dispersion at *t* = 11.11*s* (after 400 cilia revolutions), we calculate Ripley’s K function *K*(*r*) [36]. The k function above the reference line of random distribution indicates particle clustering and below indicates dispersion. We see the massless particles cluster near the cilia tip (*z* = 7*µm*) up to around 30*µm*; For *r >* 30*µm*, the particles show a dispersed pattern (Figure 8a). The size of a swirl, as seen in the bottom row of Figure 5, is approximately 30*µm*, suggesting that particles cluster within the swirl and disperse outside a swirl. When all the particles are considered, the K function suggests that overall particles show a dispersed pattern at *t* = 11.11*s*.

**Fig 8.**
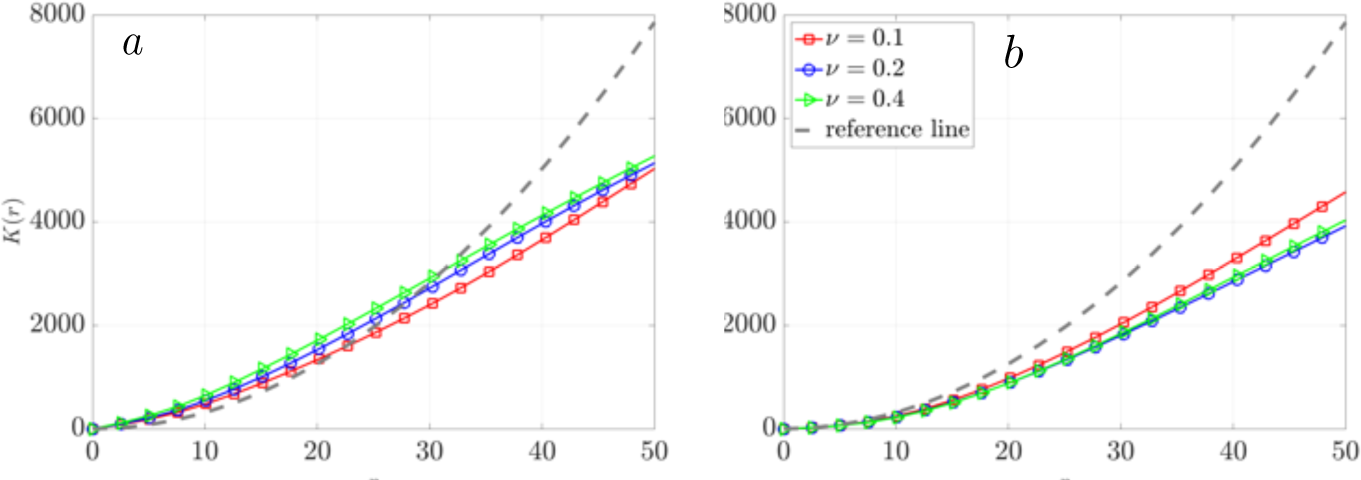
Ripley’s K function for particles at three ciliary densities *ν* = 0.1 (red), *ν* = 0.2 (blue), and *ν* = 0.4 (green) at *t* = 11.11s. (a) Particles initially located at a height of *z* = 7*µ*m. (b) All particles within the vertical range of *z* = [0, 10]*µ*m. A gray dashed reference line indicates a random distribution.

### Net transport and dispersion in a single cilia cluster

Next, we examine in detail how increasing ciliary density enhances both transport and mixing. To begin, we analyze the particle movement with a single cluster of 3 × 3 evenly spaced cilia, as illustrated in Fig. 3(a).

Over one revolution, the velocity field induced by a single cluster of cilia, as viewed from the top (Figure 9 a-d), shows a robust and uniform velocity field in four directions: upwards, leftwards, downwards, and rightwards, corresponding to the sweeping motion of the cilia. This uniform velocity pattern extends beyond the cluster boundaries and gradually diminishes radially outward. In the side view (Figs. 9(e-h)), the velocity field from the cell surface to above the cilia tip provides a more dynamic perspective. Near the cell surface, at the base of the cilia, velocity approaches zero. The velocity magnitude peaks between height of 5*µ*m to 10*µ*m near the cilia tip. During the forward stroke, the tip of the cilia is around *z* = 7*µ*m. Interestingly, when the azimuthal angle is *ϕ* = *π* (Fig. 9(g)), the velocity field appears less oriented despite its apparent uniformity from the top view (Fig. 9(c)). This discrepancy arises because the velocity vectors nearly align perpendicular to the plane of the paper at *ϕ* = *π*, coinciding with the point where the cilia begin to shorten during the backward stroke, as detailed in Equation (2).

**Fig 9.**
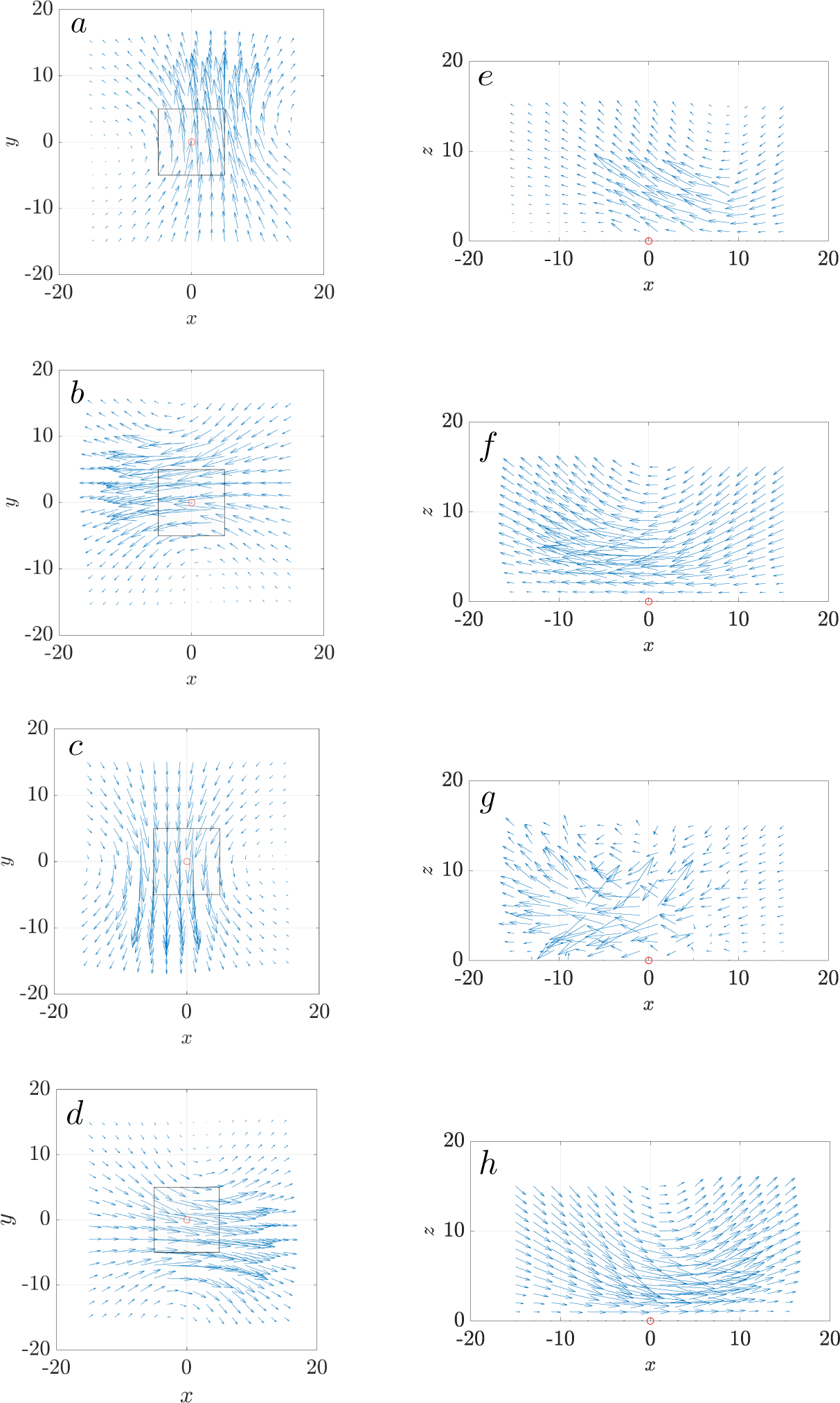
Velocity field induced by one cluster of cilia, shown from the top view at *z* = 5*µ*m in (a)-(d) and from the side view at *y* = 0*µ*m in (e)-(h). The panels correspond to the instantaneous azimuthal angles of the cilia as follows: *ϕ* = 0 (a, e), *ϕ* = *π/*2 (b, f), *ϕ* = *π* (c, g), and *ϕ* = 3*π/*2 (d, h). The black box marks the boundary of the cilia cluster, with the cluster center located at (0, 0) (red circle). All cilia beat synchronously with an initial phase angle of *ϕ*_0_ = 0.

Figure 10 shows the distribution of passive particles from both top and side views at four time points: *t* = 0*s*, 1.38*s*, 2.78*s*, and 11.11*s*, corresponding to 0, 50, 100, and 400 cilia revolutions, respectively. The particles are color-coded by height (*z*) as blue, red, and black. From the top view, a single counterclockwise swirl emerges as early as *t* = 1.38, after only 50 cilia revolutions. The swirl grows in time, extending beyond the cilia cluster boundary. As expected, the bottom (black) particles exhibit minimal motion, the mid-height (red) particles move less than the top (blue) particles. Overall, we see a net transport of particles to the lower left, in the negative *x* and *y* directions.

**Fig 10.**
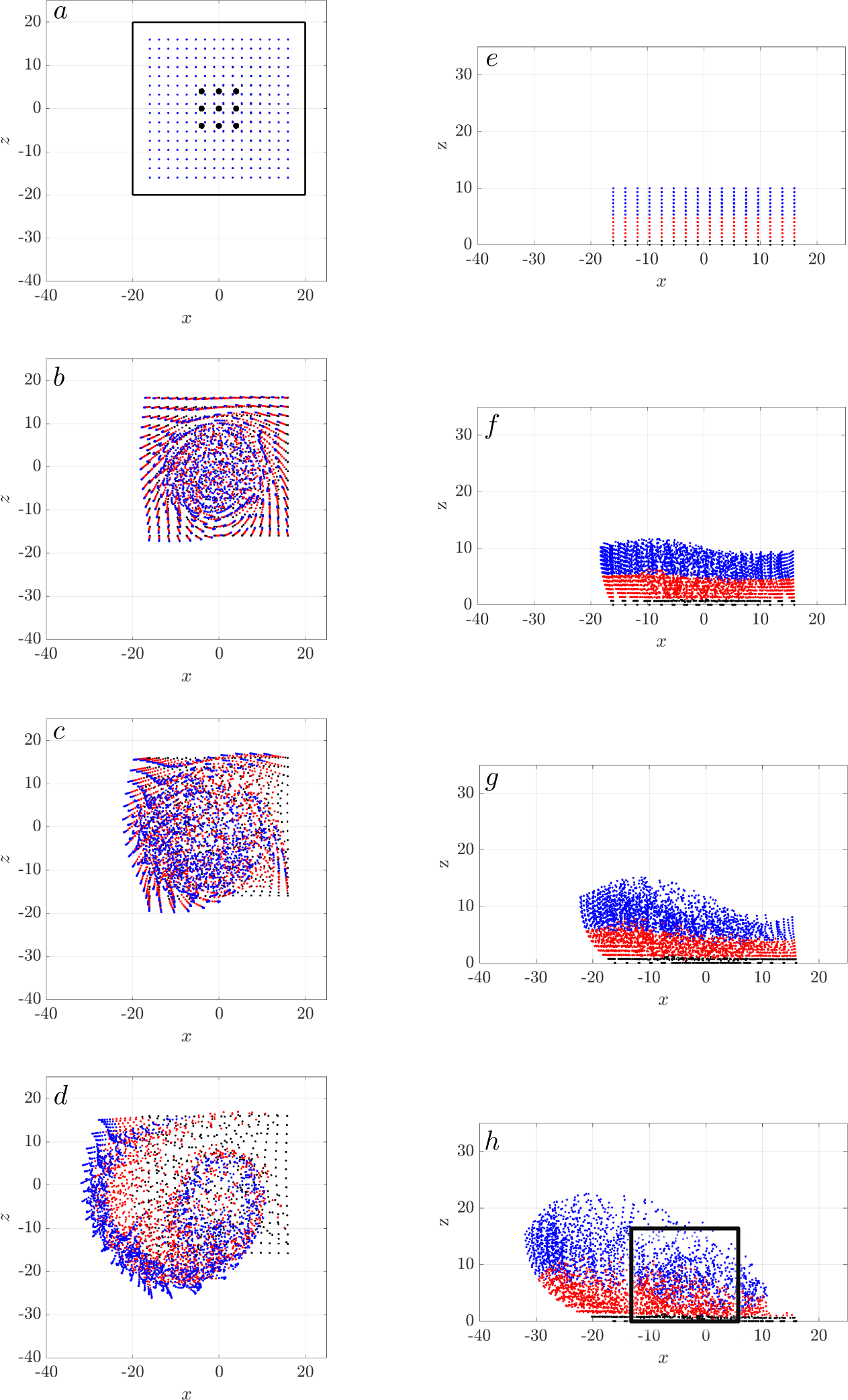
Swirl motion induced by a single cilia cluster: Distribution of passive particles visualized in top view (left column: (a)-(d)) and side view (right column: (e)-(h)). Time progresses from top to bottom, with snapshots at *t* = 0*s* in (a) and (e); *t* = 1.38*s* in (b) and (f); *t* = 2.78*s* in (c) and (g); and *t* = 11.11*s* in (d) and (h). These time points correspond to 0, 50, 100, and 400 cilia revolutions, respectively, with all cilia beat synchronously. The black box in (a) outlines the boundary ofthe cilia cluster boundary, and the blue dots indicate the positions of the 9 individual cilia. In (e), particles are color-coded by their height (*z*): blue, red, and black represent different elevation range.

From the side view, the vertical mixing of the blue and red particle layers is evident. The mixing pattern is non-uniform, as indicated in the black box around *x* ∈ [−10, 2]*µ*m in Fig. 10(h), where the blue and red particles intermingle more extensively than in regions outside the black box. Despite this non-uniformity, all particles exhibit an upward movement trend in the *z* direction.

The MSD value at *z* = 1*µ*m is significantly smaller than those at higher heights, as the mucus flow is minimal near the bottom of the cilia, close to the cell surface. For particles at higher layers (*z >* 1*µ*m), the MSD plots collapse into a single curve with a slope close to 1, followed by a tail with a slope greater than 1. This behavior suggests that, over a short time, particle spreading resembles simple diffusion even in this deterministic model. However, over longer timescales (*t >* 60*s*), directional particle transport becomes increasingly evidence (Fig. 11).

**Fig 11.**
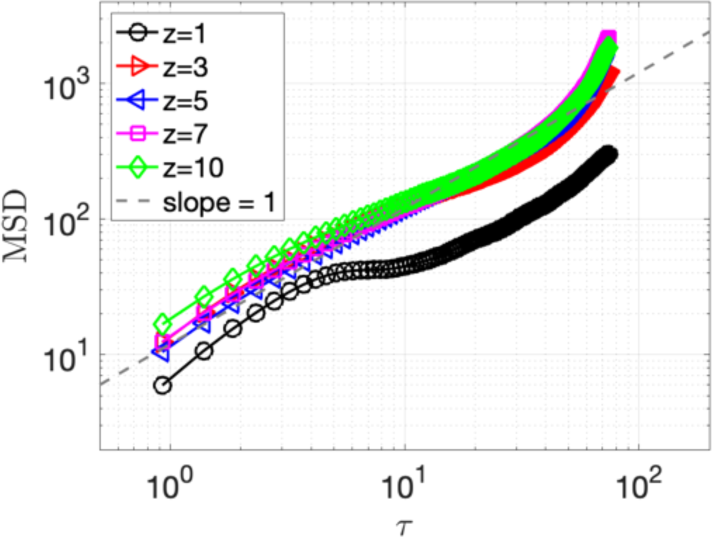
Mean squared Displacement (MSD) plots for the massless particles in mucus flow driven by a single synchronized cilia cluster, shown at different initial heights: *z* = 1*µ*m, 3*µm*, 5*µ*m, 7*µ*m and 10*µ*m. A gray reference line has a slope of 1. Total simulation time is 5 days.

We further calculate the Ripley’s K function to quantify the spatial distribution of the particles at *t* = 11.11*s*. Fig. 12 shows *K*(*r*) for particles at different heights *z*(*µ*m), with the dashed reference line representing the completely random distribution. Particles at *z* = 1 exhibit minimal movement, resulting in a curve (black) that falls below the reference line. Particles at all *z >* 1(*µ*m) display significant clustering, as their curves lie above the reference line for *r* ≤ 6(*µ*m). For *r >* 6*µm*, particles show dispersion as their curves fall below the reference line. This pattern of short range clustering and longer range dispersion is associated with the swirls observed in Fig. 10. Clustering is most pronounced at *z* = 5 and *z* = 7, near the cilia tips.

**Fig 12.**
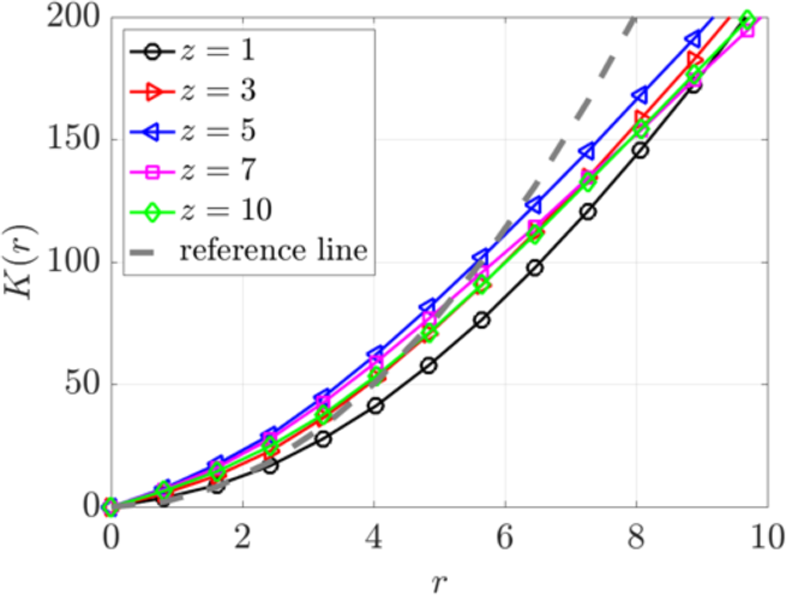
Ripley’s K function for particles at various heights (*z* = 1, 3, 5, 7, 10)*µ*m at *t* = 11.11*s* for a single cilia cluster. A dashed reference line indicates a random distribution.

### Maximal transport in array of synchronized cilia clusters with optimal spacing

As each cilia cluster generates a swirl that shows a net transport and mixing, how do cilia clusters interact with each other? Next, we examine the transport and mixing with an array of three synchronized cilia clusters. We vary the spacing between clusters,*D_c_*, from 3*D* to 12*D*, with *D* = 4*µ*m, as illustrated in Fig. 3. When *D_c_* = 3*D*, these three cluster boxes are adjacent to each other, forming one continuous cluster three times as long. Consequently, this arrangement establishes an extended platform of the velocity field, leading to a more pronounced and uniform motion in the *x* direction (Figure 13).

**Fig 13.**
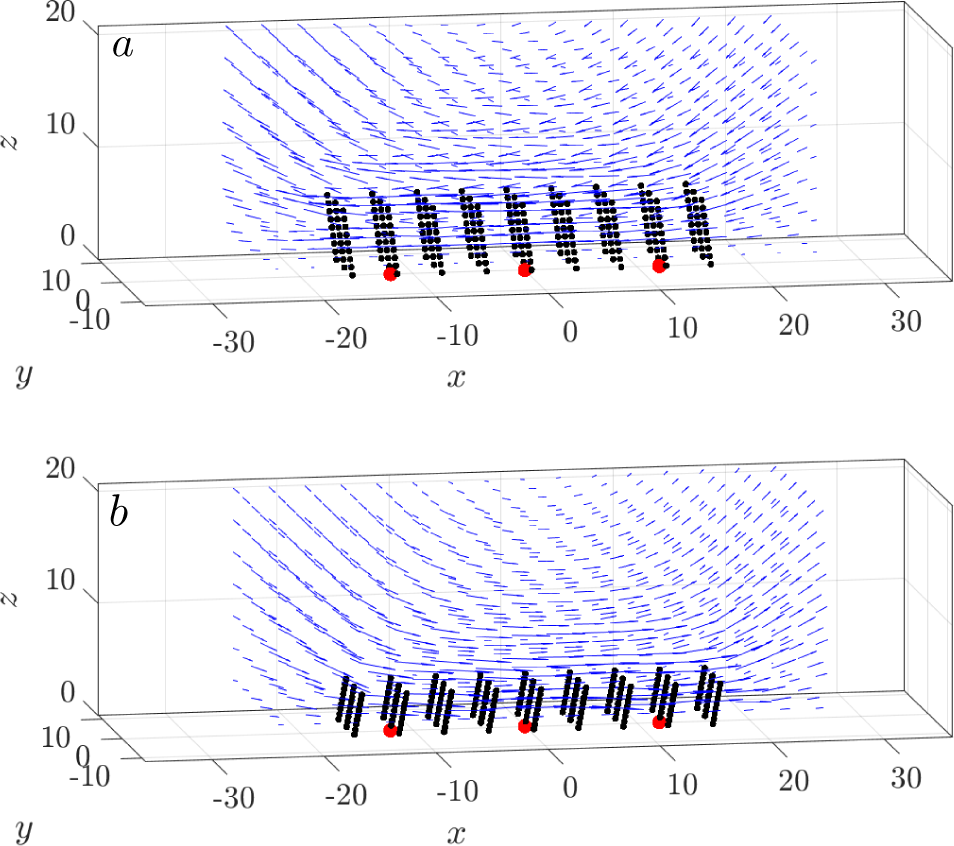
Uniform horizontal motion in one large cluster form by placing three cilia clusters next to each other, *D_c_* = 3*D*. The 3-dimensional velocity field and the orientation of all cilia at*ϕ* = *π/*2 (a), and *ϕ* = 3*π/*2 (b). All cilia beat synchronously. The red dots indicate the cluster centers.

We color the passive, massless particles based on their initial *x* positions to visualize their motion induced by three adjacent cilia clusters. The top-view plots of the particles (Fig. 14 a-d) show the formation of a counterclockwise swirl, similar to the pattern observed with a single cilia cluster (Fig. 10 a-d) but on a larger scale. Over time, the swirl expands and travels toward the lower-left direction, accompanied by significant horizontal mixing as particles of different colors intermix.

**Fig 14.**
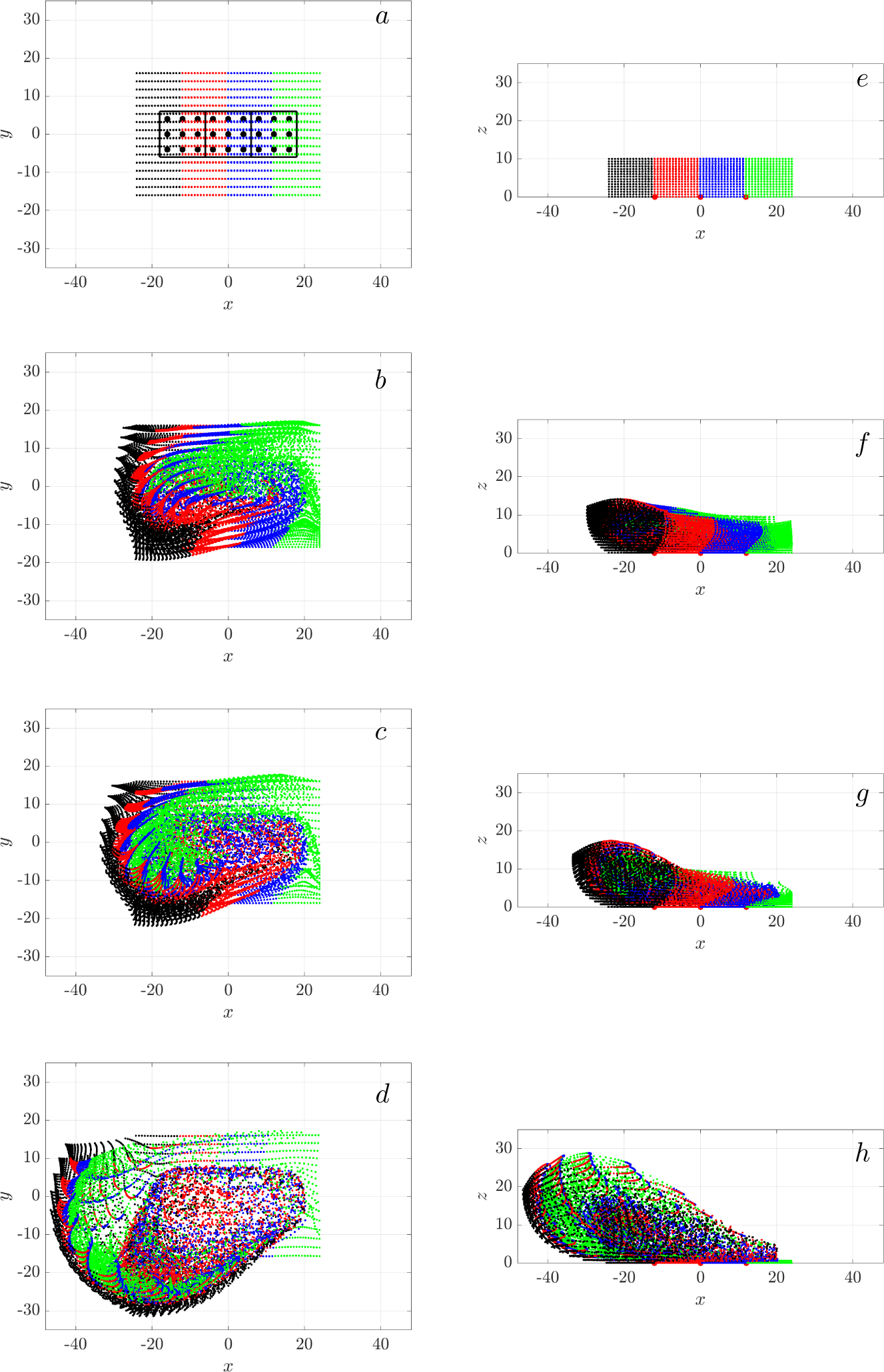
Transport and mixing of particles in a cluster of three adjacent cilia clusters, separated by *D_c_* = 3*D*. The left column (a-d) shows a top view, and the right column (e-h) shows a side view. From top to bottom, sequential snapshots represent time points: *t* = 0*s* (a,e), *t* = 1.38*s* (b,f), *t* = 2.78*s* (c,g), and *t* = 11.11*s* (d,h), corresponding to 0, 50, 100, and 400 cilia revolutions, respectively. All cilia movesynchronously. Each cluster is marked by black boxes. Particle colors indicate their initial positions along the *x*-axis, illustrating the extend of horizontal transport and mixing.

In contrast, the side-view plots (Figs. 14 e-h) show a predominant movement of green particles, initially positioned at the far right, migrating towards the left. Remarkably, particles from all four color groups exhibit significant mixing in the *x* direction, as evidenced by the presence of all colors near the left boundary (Fig. 14h). This pronounced mixing and transport result from the close proximity of the three cilia clusters, which collectively behave like a single large cluster, facilitating efficient mixing and transport.

At the opposite end of the spacing spectrum, with the cluster spacing set to *D_c_* = 12*D*, the clusters are sufficiently separated to each generate individual swirls (Fig. 15). Over time, the swirls expand into adjacent areas, initiating particle mixing at the interfaces. The particles move upwards. But unlike the closely spaced clusters previous described, the majority of the four particle colors remain largely distinct along the *x* direction, indicating limited mixing between them.

**Fig 15.**
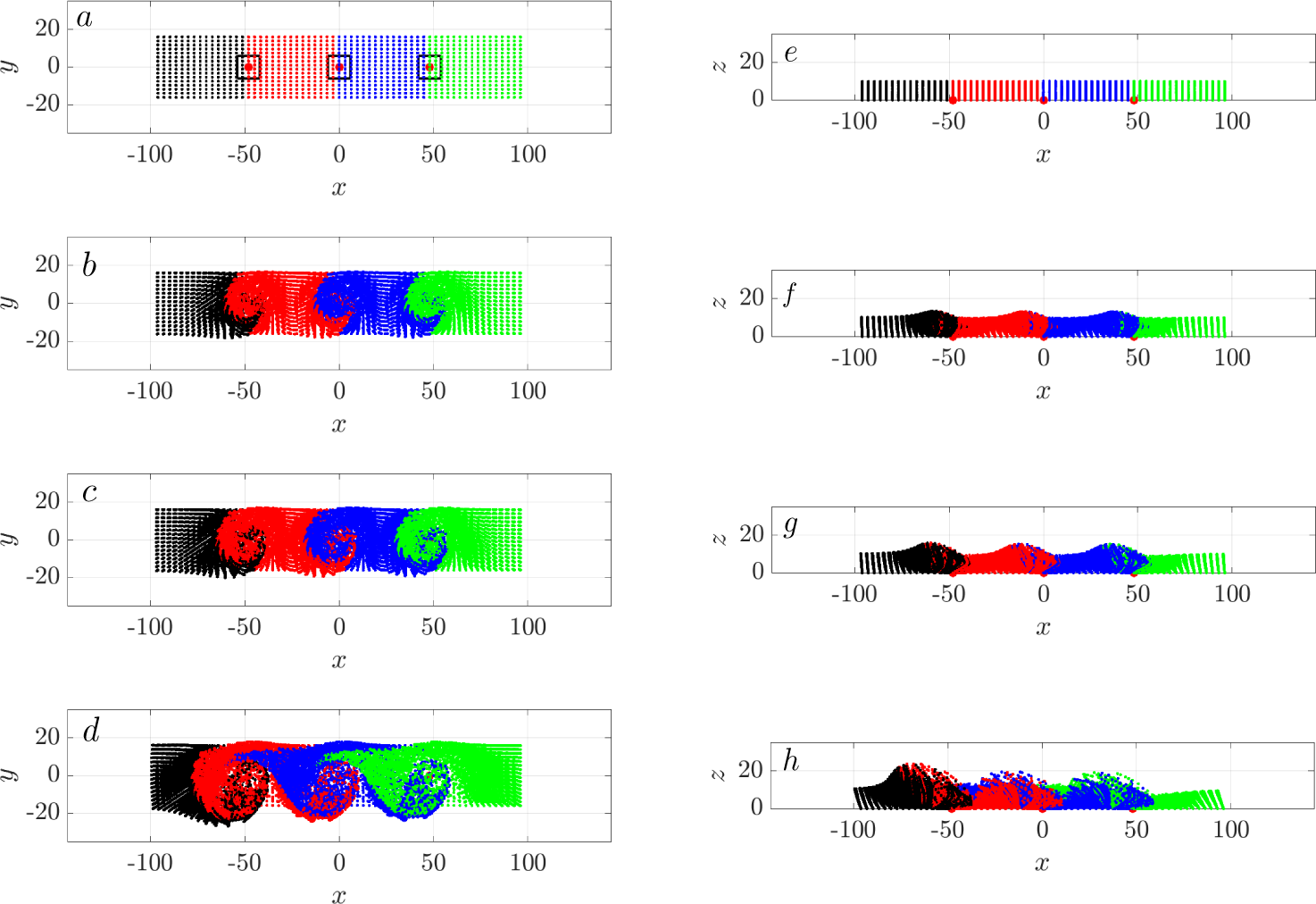
Swirls from three widely separated cilia clusters, spaced at *D_c_* = 12*D*. The left column shows the top view (a-d) and the right column shows the side view (e)-(h). From top to bottom, sequential snapshots represent time points: *t* = 0*s* (a,e), *t* = 1.38*s* (b,f), *t* = 2.78*s* (c,g), and *t* = 11.11*s* (d,h), corresponding to 0, 50, 100, and 400 cilia revolutions, respectively. All cilia beat synchronously. Cluster centers are marked with red dots. Particle colors represent their initial positions along the *x*-axis, illustrating the dynamics of mixing and transport.

It is now evident that cluster spacing significantly influences particle transport dynamics. We see limited net particle transport at both extreme cluster spacings of *D_c_* = 3*D* and 12*D*. To determine if there is an optimal cluster spacing that maximizes net transport, we vary *D_c_* between 3*D* and 12*D* and analyze the net transport as measured by the center of mass of all the particles. As shown in Figure 16(a), the maximal transport in *x*-direction (along the airway) occurs at an intermediate spacing of *D_c_*= 5*D*. Interestingly, both transverse (*y*−direction) and vertical transport (*z*−direction) decrease monotonically with increasing inter-cluster spacing, with the maximum transverse and vertical transport observed when the clusters are next to each other.

**Fig 16.**
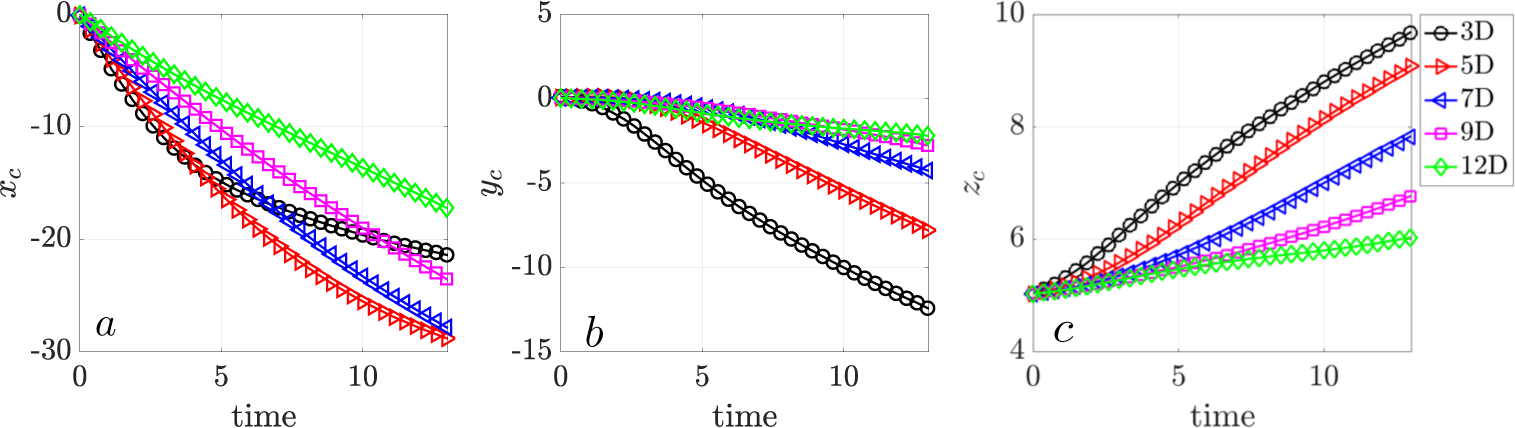
Maximal transport at intermediate cluster spacing. The trajectories of the center of mass (CoM) for all particles for three cilia clusters, in the (a) *x*-direction, (b) *y*-direction, and (c)*z*-direction. All cilia beat synchronously.

The mean square displacement (MSD) for increasing cluster spacing *D_c_* between three synchronized cilia clusters shows distinct particle motion characteristics (Figs. 17). The slopes of the MSD plots in logarithmic scale are all greater than 1, indicating particles are transported instead of staying in a confined region. Particles at *D_c_* = 5*D* and *D_c_*= 7*D* generally travel further, as their MSD slopes exceed those at *D_c_*= 3*D* and *D_c_* = 12*D*. This observation reflects the role of hydrodynamic interactions in particle transport. At *D_c_* = 3*D*, the clusters are too closely spaced, limiting effective particle displacement. Conversely, at *D_c_* = 12*D*, the clusters are too far apart to interact effectively. The intermediate spacing of *D_c_*= 5*D,* 7*D* achieves a balance, leveraging hydrodynamics to enhance particle transport. Furthermore, the largest MSD slope values occur at *z* = 5 and 7*µ*m, corresponding to regions near the cilia tips. These regions exhibit the highest velocity, further emphasizing the critical role of cilia-tip dynamics in driving effective particle transport.

**Fig 17.**
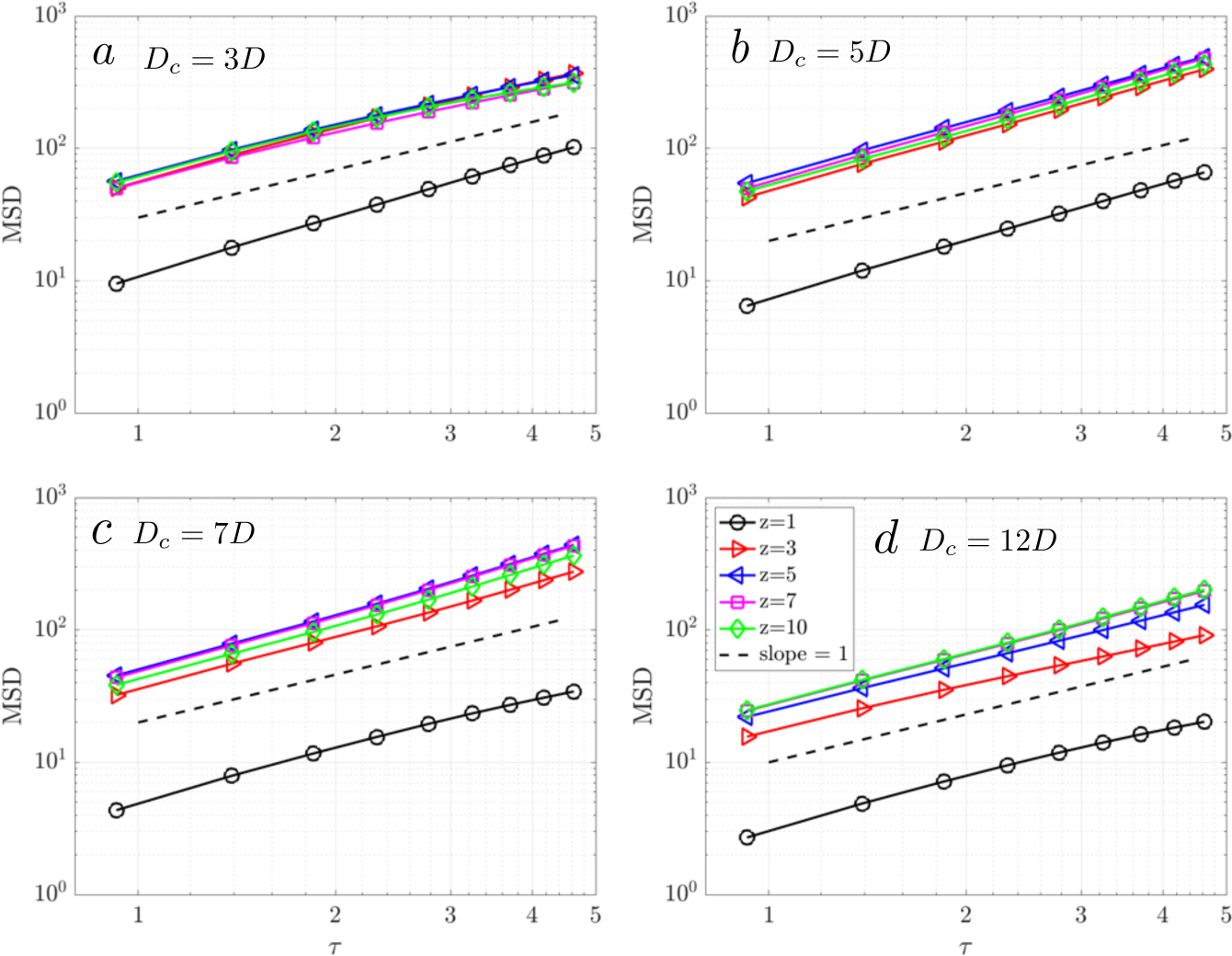
Mean square displacement (*MSD*) of particles at various initial heights (*z* = 1, 3, 5, 7, 10 *µ*m) across different cilia spacings for three cilia clusters: (a) *D_c_* = 3*D*, (b) *D_c_* = 5*D*, (c) *D_c_* = 7*D*, (d) *D_c_* = 12*D*. All cilia beat synchronously.

Figure 18 shows Ripley’s K function analysis for particles in three cilia clusters with varying cluster spacing from *D_c_* = 3*D* to 12*D*. We focus on particles near the cilia tip at a height of *z* = 7*µ*m. The results show evident short-range particle aggregation and long-range dispersion across all spacings. The sizes of these aggregates correlate with the swirl sizes and increase with increasing *D_c_*, reaching a plateau for spacing greater than *D_c_*= 10*D*.

**Fig 18.**
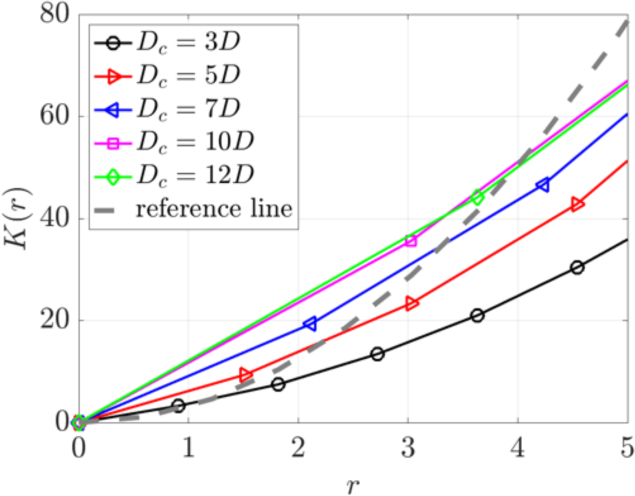
Ripley’s K function analysis of particle distributions in three cilia clusters with varying spacing from *D_c_* = 3*D* to *D_c_* = 12*D*. This analysis focused on particle positions at a height of *z* = 7*µ*m. A gray dashed reference line represents a random distribution.

Taken together, the swirls generated by individual cilia clusters interact hydrodynamically, enhancing both directional transport and mixing. Importantly, there is an optimal spacing between the cilia clusters that maximizes the speed of directional transport when all cilia beat synchronously.

### Metachronal wave inhibits transport

Thus far we have focused on synchronous cilia beating. On the airway surface with many cilia, the actuating organelles can coordinate with each other and collectively beat in the form of metachronal waves, where neighboring cilia beat sequentially (i.e., with a phase lag) rather than synchronously [37]. Metachronal waves in mammals can move 4–8 times faster than mucociliary transport. In the trachea, these waves can move at speeds of 6–20 mm per minute [38]. The cilia metachrony is characterized based on the phase lag between each other and it arises via hydrodynamics interactions [39]. The metachrony is categorized into four types: antipleptic (phase lag *ϕ*_0_ ∈ [0*, π*]), symplectic (*ϕ*_0_ ∈ [*π,* 2*π*]), synchronized (*ϕ*_0_ = 0), and standing wave (*ϕ*_0_ = ±*π*).

Here, we assume that the metachronal wave is generated by the actuation of organelles rather than emerging from hydrodynamic interactions between beating cilia. We study the effect of metachronal wave on mucociliary transport by imposing the phase lags directly between cilia. We focus on the scenario of three clusters with a cluster spacing of *D_c_* = 3*D*, varying the phase lag *ϕ*_0_ between cilia. Results for other values of *D_c_* exhibit similar trends and are therefore not shown. The phase lag is applied to the cilia arrays in the *x*-direction. Specifically, each 3 × 3 cilia cluster is organized into 3 columns based on their *x* locations. Each column begins with an equal increment of *ϕ*_0_, while maintaining synchronicity among the cilia within the same column.

With metachronal waves, the distribution of particles at *t* = 11.11*s*, corresponding to 400 cilia revolutions, show reduced net transport and increased mixing (Fig. 19). For *ϕ*_0_ /= 0, particles aggregate into a large swirl. As the phase lag increases, more particles become confined to the central region, where they are well-mixed, and the net transport of particles in the negative *x* and *y* directions progressively diminishes. At *ϕ*_0_ = 2*π/*3, all particles are effectively trapped, and the swirl transformes into an ellipse. This occurs because the phase lag is commensurate with the cilia array’s periodicty: after 3 columns, the phase lag totals 2*π*, equivalent to no phase lag between clusters. Thus, the metachronal wave suppresses directional transport in the *x* − *y* plane within a large cilia cluster.

**Fig 19.**
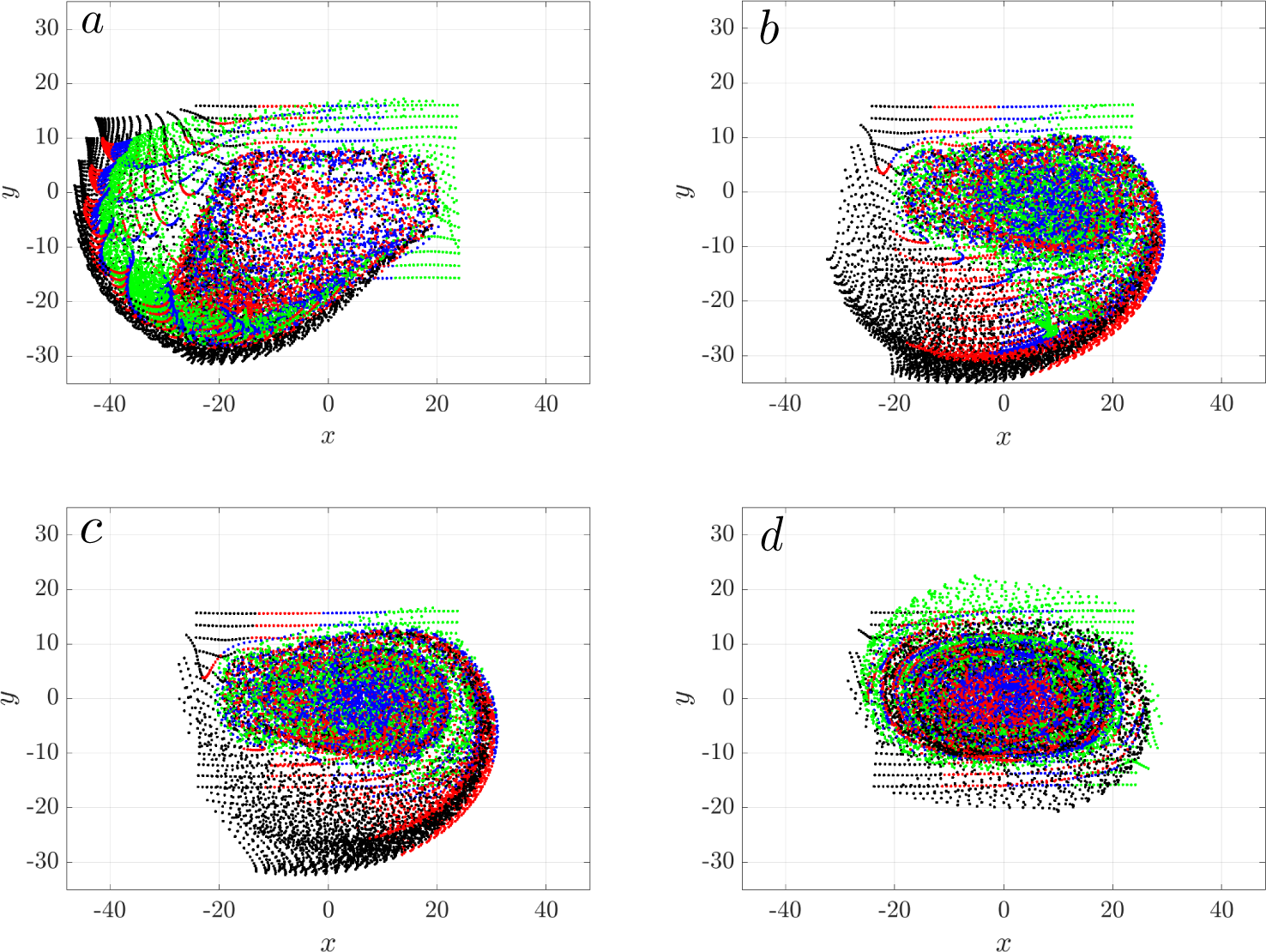
Metachronal wave reduces directional transport in three connected cilia clusters with spacing *D_c_* = 3*D*. Snapshot of massless particles at *t* = 11.11*s* are shown for varying phase shifts applied along the *x* direction. (a) *ϕ*_0_ = 0, (b) *ϕ*_0_ = *π/*4, (c) *ϕ*_0_ = *π/*3, (d) *ϕ*_0_ = 2*π/*3. At *ϕ*_0_ = 2*π/*3, the three columns exhibit a metachronal wave with a wavelength of 2*π*. Particles are color-coded based on their initial positions in the *x* direction at *t* = 0, same as in Fig. 14(a).

We analyze the CoM trajectories of particles for varying phase lags *ϕ*_0_ ∈ [0, 2*π/*3] (Figure 20). As *ϕ*_0_ increases from 0 to *π/*3, the direction of transport along the *x*-axis, denoted by *x_c_*, shifts from negative to positive, indicating a reversal in transport direction. However, the magnitude of *x_c_* diminishes as *ϕ*_0_ increases, approaching zero as *ϕ*_0_ nears 2*π/*3. Notably, at *ϕ*_0_ = *π/*4 and *π/*3, the metachronal wave reverses the particle transport direction. In terms of transverse transport (*y_c_*), a phase lag of *ϕ*_0_ = 2*π/*3 results in minimal change. At *ϕ*_0_ = *π/*4 and *π/*3, *y_c_* initially decreases and then stabilizes, whereas for *ϕ*_0_ = 0 and *π/*12, *y_c_* shows a continual decrease, albeit with smaller magnitudes than those seen in *x_c_*. No reverse transport in the positive *y*-direction is observed.

**Fig 20.**
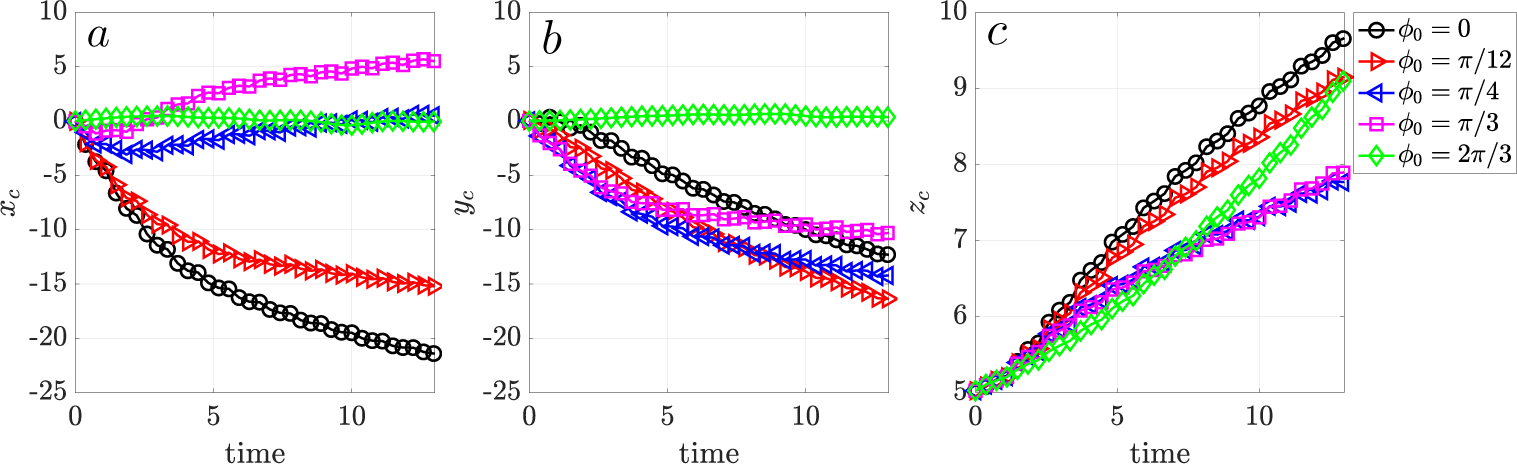
Transport of particles in an array of three connected cilia cluster with metachronal waves, shown for phase lags *ϕ*_0_ = 0, *π/*12, *π/*4, *π/*3, 2*π/*3. The center of mass (CoM) of all particles is plotted for (a) the *x*-direction, (b) the *y*-direction, and (c) the *z*-direction.

Interestingly, metachrony significantly enhances vertical transport *z_c_* away from the cell surface, moving in the positive *z*-direction. As shown in Figure 20 (c), *z_c_* increases across all values of *ϕ*_0_. The trajectory for *ϕ*_0_ = 2*π/*3 closely approximates a linear increase, whereas those for other phase values initially rise more steeply than linearly before slowing down. This pattern occurs because metachrony generates rotating vortex flows that drive particles upward. This dual influence of metachronal waves on particle transport underscores the intricate interaction between ciliary coordination and fluid dynamics within the mucociliary system.

## Discussion

The ciliated cells of the airway epithelium play a crucial role in facilitating mucociliary clearance, which helps clear the airway of pathogens and other particles. In this study, we use computational tools to simulate cilia activity and mucus transport on a small tissue scale, and examine the transport and mixing of tiny particles on a tissue patch, an array of three cilia clusters, and then a single cluster.

The size of a tissue patch is comparable to that of human bronchial culture in a recent experimental study [5]. In this case, beating cilia induce swirling flow patterns, similar to those observed in human bronchial tissue experiments [5, 40]. We found that transport is indeed enhanced at higher ciliary densities. However, interestingly, the size of particle clustering or swirling patterns was found to be independent of the ciliated density. This contrasts with findings from Gsell et al. (2020) [8]. The circular pattern induced by a patch of cilia clusters are also observed in the experimental study of Sears et al. (2015) [41] using primary human airway epithelial cells and referred to as mucus hurricane. Such observations have been instrumental in investigating the pathogenesis of cystic fibrosis [42]. These swirling patterns and their implications in mucus transport dynamics highlight the intricate interplay between ciliary motion, fluid dynamics, and respiratory health.

While many studies have investigated the formation of metachronal waves and their role in either facilitating or hindering the efficiency of fluid pumping [25, 27, 31, 43, 44], the results remains inconclusive. For example, Gauger et al. [43] modeled cilia with super-paramagnetic particles in an external magnetic field and found that antipleptic metachrony was more efficient for transport with a particular cilia spacing. Wollin and Stark [25] modeled cilia as chains of beads and found transient synchronization in a bulk fluid. Elgeti and Gompper [31] represented cilia as semi-flexible rods with active bending forces, identified both symplectic and laeoplectic (perpendicular to the power-stroke direction) metachrony, and noted a 10-fold increase in propulsion efficiency compared to synchronous beating. Our study prescribes antipleptic metachrony, as the phase lag is within [0*, π*]. Interestingly, we observed that a phase lag greater than *π/*4 among cilia induces reverse transport in the *x*-direction within the mucus layer. This finding is consistent with the results from Chateau et al. [44], who reported an inhibitory effect of metachrony on transport.

Understanding the characteristics of particle transport and mixing within the mucociliary system is crucial for comprehending disease pathophysiology and developing effective treatment strategies. By precisely prescribing the motion of 3D cilia, we can isolate and study the effects of various parameters, thereby taking a fundamental step toward understanding the transmission of particles such as bacteria, viruses, and inhaled drugs. The insights gained from our current work provide valuable knowledge about particle transport and mixing in mucus fluid flow. This knowledge can contribute to the development of novel therapies, diagnostic techniques, and medical devices aimed at addressing respiratory diseases and improving overall respiratory health. Through continued research and innovation in this field, we can advance our understanding of mucociliary dynamics and pave the way for more targeted and effective interventions in respiratory health management.

## Acknowledgment

YJ acknowledges the support from the Frady Whipple Professorship. We thank the Advanced Research Computing Technology & Innovation Core (ARCTIC) team at GSU for support.

